# Concurrent origins of the genetic code and the homochirality of life, and the origin and evolution of biodiversity. Part I: Observations and explanations

**DOI:** 10.1101/017962

**Authors:** Dirson Jian Li

## Abstract

The post-genomic era has brought opportunities to bridge traditionally separate fields on early history of life. New methods promote a deeper understanding of the origin of biodiversity. Relative stabilities of base triplexes are able to regulate base substitutions in triplex DNAs. We constructed a roadmap based on such a regulation to explain concurrent origins of the genetic code and the homochirality of life. Based on the recruitment order of codons in the roadmap and the complete genome sequences, we reconstructed the three-domain tree of life. The Phanerozoic biodiversity curve has been reconstructed based on genomic, climatic and eustatic data; this result supports tectonic cause of mass extinctions. Our results indicate that chirality played a crucial role in the origin and evolution of life. Here is Part I of my two-part series paper; technical details are in Part II of this paper (see “Concurrent origins of the genetic code and the homochirality of life, and the origin and evolution of biodiversity. Part II: Technical appendix” on bioRxiv).

---

The origins of the genetic code, the homochirality of life and the biodiversity are among the key problems on the origin of life. These fundamental problems are traditionally treated separately, for each of which numerous explanations have been proposed. For example, frozen accident, error minimisation, stereochemical interaction, amino acid biosynthesis and expanding codons have been suggested to explain the origin of the genetic code [1] [2] [3] [4]; biochemical, geochemical or interstellar processes may account for the origin of the homochirality of life [5] [6] [7]; and the three-domain tree and the eocyte tree are candidates for the tree of life [8] [9]. Debates exist and remain unresolved. The situation has not taken a good turn due to lack of criteria to judge among these explanations.

The purpose of this paper is to explore these problems based on biological and geological data. The above events at the beginning of life were interrelated. Considering base substitutions in triplex DNA, a roadmap for the evolution of the genetic code is proposed. The universal genetic code are achieved by the roadmap considering that the stabilities of base triplexes tend to increase in the substitutions. The roadmap is chiral, either right-handed or left-handed. This consequently indicates that the homochirality of life resulted from the competition between the two chiral roadmaps.

The roadmap brings new insight into the origin of the three domains of life. In section 2, we show that the three-domain tree of life can be reconstructed based on the statistical differences of complete genome sequences between certain groups of codons in the roadmap, which indicates that the root of the tree of life was determined by the evolution of the genetic code. Thus, the origins of the genetic code, the homochirality and the three domains altogether have been explained by the roadmap theory. Explaining these problems together is easier than explaining them separately, because borrowed ideas from each other enhance the explanations themselves.

In section 3, we show that the trend and the fluctuations of the Phanerozoic biodiversity curve [10] can be explained respectively by the genome size evolution and by the climate changes and sea level fluctuations. The five mass extinctions are discerned in our reconstructed biodiversity curve. And, the declining origination rate and extinction rate through the Phanerozoic are explained. We reveal that the homochirality of life determines the direction of the evolution of life at the sequence level, which guarantees the adaptation of life to the changing Earth system.

## 1 Concurrent origins of the genetic code and the homochirality of life

A roadmap for the evolution of the genetic code (the roadmap for short, Fig 1A) has been constructed based on the relative stabilities of base triplexes [11] [12] in the base substitutions in triplex DNA, as shown below.

**Fig 1.**
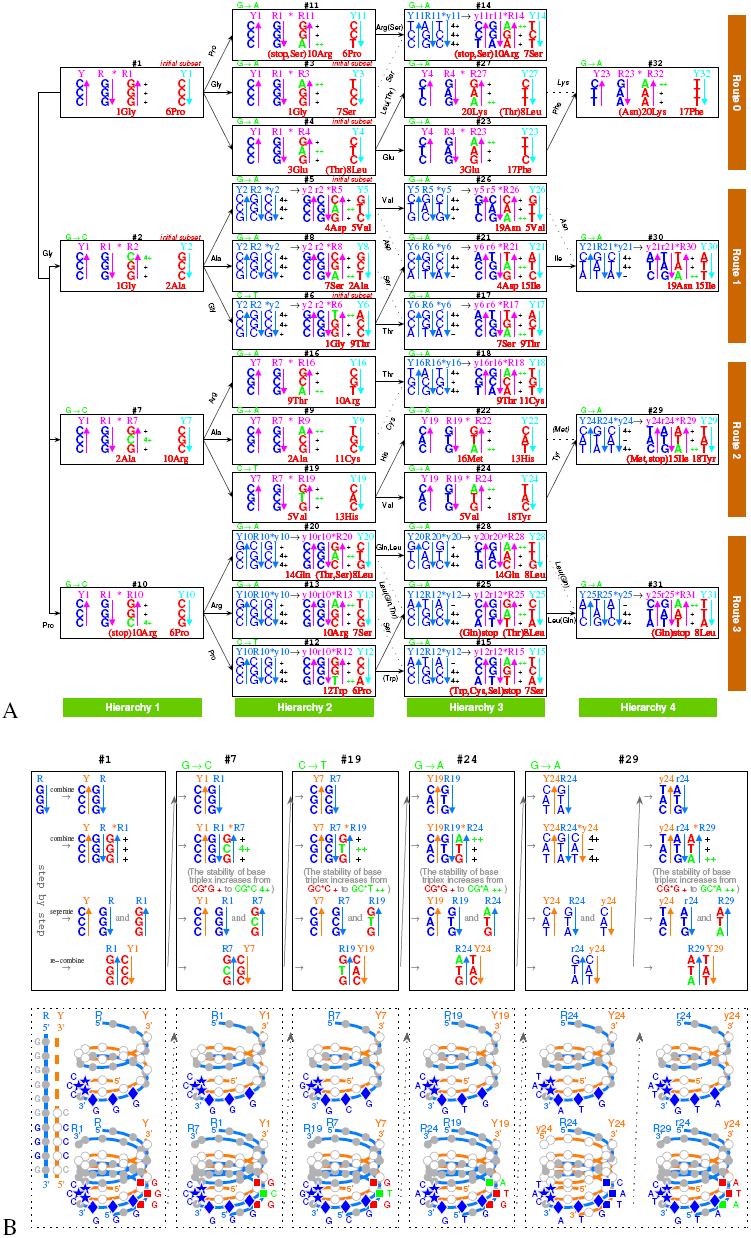
Explanation of the origin of the genetic code. **A**, The roadmap for the evolution of the genetic code. The 64 codons formed from base substitutions in triplex DNAs are in red (see Appendix for detailed caption, same below). **B**, A detailed description of the roadmap in the right-handed triplex DNA picture, taking for example from #1 to #29.

At the beginning of the evolution of the genetic code, there existed single-stranded DNA *Poly G* and *Poly C*, which tended to form a triplex DNA [11] [13]. *Poly C* · *Poly G* * *Poly G* is a usual *YR* * *R* triplex DNA, which is formed by base triplex *CG* * *G* (Fig 1B). Since *CG* * *A* is more stable than *CG* * *G* [11] [12], the transition from *G* to *A* occurs spontaneously, by which all the codons in *Route* 0 were recruited (Fig 1A). Since *CG* * *C* is more stable than *CG* * *G*, the transversion from *G* to *C* occurs spontaneously, which blazed a new path for the recruitment of codons in *Route* 1 − 3 (#2, #7, #10 in Fig 1A). And since *GC* * *T* is more stable than *GC* * *C*, the transition from *C* to *T* also occurs spontaneously, by which the remaining codons in *Route* 1 − 3 were recruited (Fig 1A). Thus, all the 64 codons have been recruited following the roadmap (Fig 1). The roadmap had avoided the unstable base triplexes [11] (Fig S1a1). The choice of the genetic code was by no means random, which resulted from the increasing stabilities of base triplexes in the substitutions.

According to the roadmap, tRNAs have been obtained to transport all the amino acids respectively (Fig 2A). For each of the 20 amino acids, a pair of complementary single-stranded RNAs are generated at certain node in the roadmap, which joined and folded into a tRNA [14] whose anticodon encodes the corresponding amino acid (Fig S2a3). Such tRNAs help to implement the Central Dogma and to explain the codon degeneracy.

**Fig 2.**
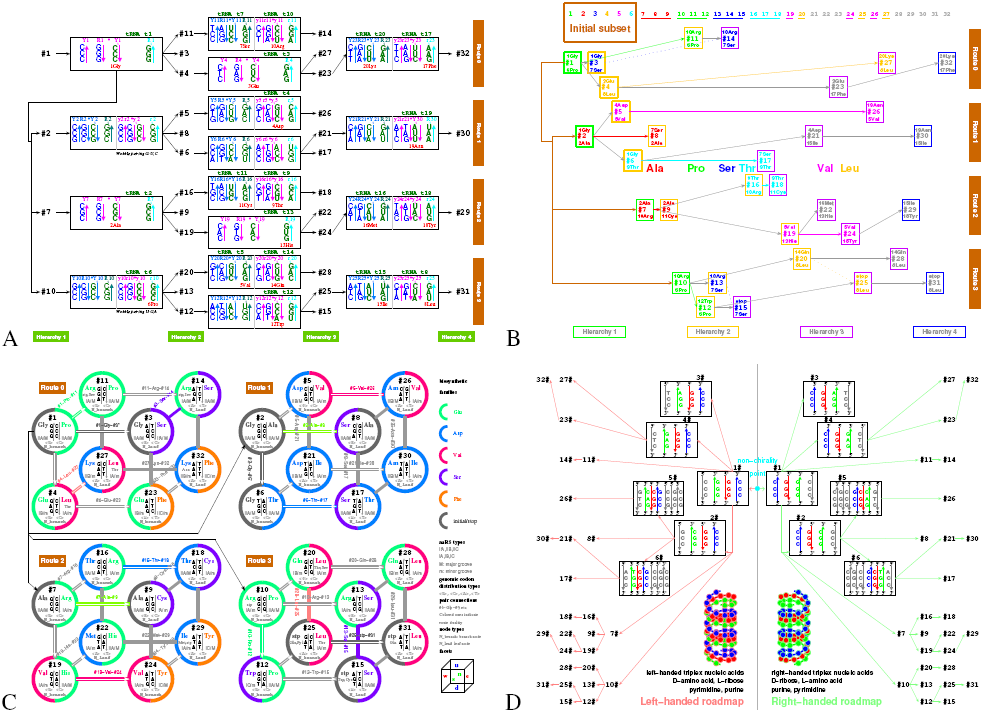
Explanations of the origin of tRNAs, codon degeneracy, and the origin of homochirality of life. **A**, The tRNAs originated along the roadmap are able to transport all the canonical amino acids. **B**, Cooperative recruitments of codons and amino acids along the roadmap. **C**, Explanation of the codon degeneracy based on the relationships among codons (pair connections and route dualities) in the roadmap. **D**, The homochirality of life originated in a winner-take-all game between the right-handed roadmap and the left-handed roadmap.

The recruitment order from #1 to #32 for the codon pairs and the recruitment order from *No.*1 to *No.*20 for the amino acids were intricately well organised and coherent (see Appendix), as shown in the roadmap (Fig 1A, 2B). In the initiation stage of the evolution of the genetic code, all the codons in the initial subset (namely #1 to #6 codon pairs) encode Phase *I* amino acids [15] respectively (Fig 2B Initial subset). In its expansion stage, each of the 6 codon pairs in the initial subset brought 3 additional codon pairs, by which the Phase *II* amino acids [15] were recruited (Fig 2B). In its ending stage, the remaining codons were recruited (Fig 2B). The three stop codons were recruited along *Route* 3, which is the last route in the roadmap (Fig 1A).

The codon degeneracy can be explained by the recruitments of codons and amino acids in the roadmap (Fig 1A, 2C). According to the order of base substitutions, the roadmap can be divided into four routes *Route* 0 − 3 from top to bottom or four hierarchies *Hierarchy* 1 − 4 from left to right (Fig 1A, 2C). Pair connections and route dualities in the roadmap (see Appendix) assisted the expansion of the genetic code as well as the assignment of codons (Fig 1A, 2B, 2C). According to the positions of amino acids in the roadmap (Fig 1A, 2C), the degeneracies of most early amino acids are 4 or 6 due to pair connections and route dualities, while the degeneracies of late amino acids are smaller. Non-standard codons also obey pair connections and route dualities, most of which appear in the last route *Route* 3 (Fig 1A).

Additionally, the roadmap is supported by the following observations (see Appendix). The variations of codon position *GC* contents and the stop codon usage in observations agree with the results by the roadmap (comparing Fig S2b2 with Figure 2 in [16] or Figure 5 in [17] for the former, and comparing Fig 2b3 with Figure 1 in [18] for the latter). The recruitment order of codons in the literatures [19] generally agrees with the recruitment order of codons in the roadmap. And the recruitment order of the amino acids obtained from the roadmap can be used to explain the variation trends of amino acid frequencies [20] (Fig S2b4, S2b5).

From the roadmap, the homochirality of life can be derived directly (Fig 2D). The initial environment on Earth was chirally symmetric, where there were equal amount of left- and right-handed amino acids and riboses. There would be two possible chiral roadmaps in the evolution of the genetic code: the right-handed roadmap based on right-handed triplex DNA (corresponding to D-ribose and L-amino acid) and the left-handed roadmap based on left-handed triplex DNA (corresponding to L-ribose and D-amino acid) (Fig 1, 2D). Both the right- and left-handed roadmaps share the common non-chiral purine and pyrimidine bases. Competition between the right- and left-handed roadmaps resulted in that only one chirality survived in the winner-take-all game (see Appendix). The choice of chirality, following either the right- or the left-handed roadmap, is by chance in this theory. In reality, the life on Earth followed the right-handed roadmap. The competition process lasted at least as much time as the amino acids evolved from Phase *I* to Phase *II*. The chiral symmetry breaking occurred spontaneously on Earth, which stripped the living chiral system from the non-living non-chiral system. And it led the direction of the evolution of life that is getting away from the starting non-chirality status (see Fig 2D). Homochirality of life protects the living system against the non-chiral environment in the evolution of life.

## 2 Origin of the three domains according to the roadmap and genomic codon distributions

The roadmap is directly supported by the statistical analysis of the complete genome sequences of contemporary species. The genomic codon distribution, as a useful technique, is defined for a species by the distribution of codon intervals in its complete genome sequence for the 64 codons respectively (Fig S3a3). According to the correlation coefficients among the genomic codon distributions, on one hand, a distance matrix for species is obtained by averaging the correlation coefficients among the codons, which results in a reasonable tree of life (Fig S3a5), on the other hand, a distance matrix for codons is obtained by averaging the correlation coefficients among the species, which results in a reasonable tree of codons (Fig 3A). Especially, such a tree of codons agrees with the differentiation of the four hierarchies of codons by the roadmap (comparing *Hierarchy* 1 − 4 between Fig 3A and Fig 1A).

**Fig 3.**
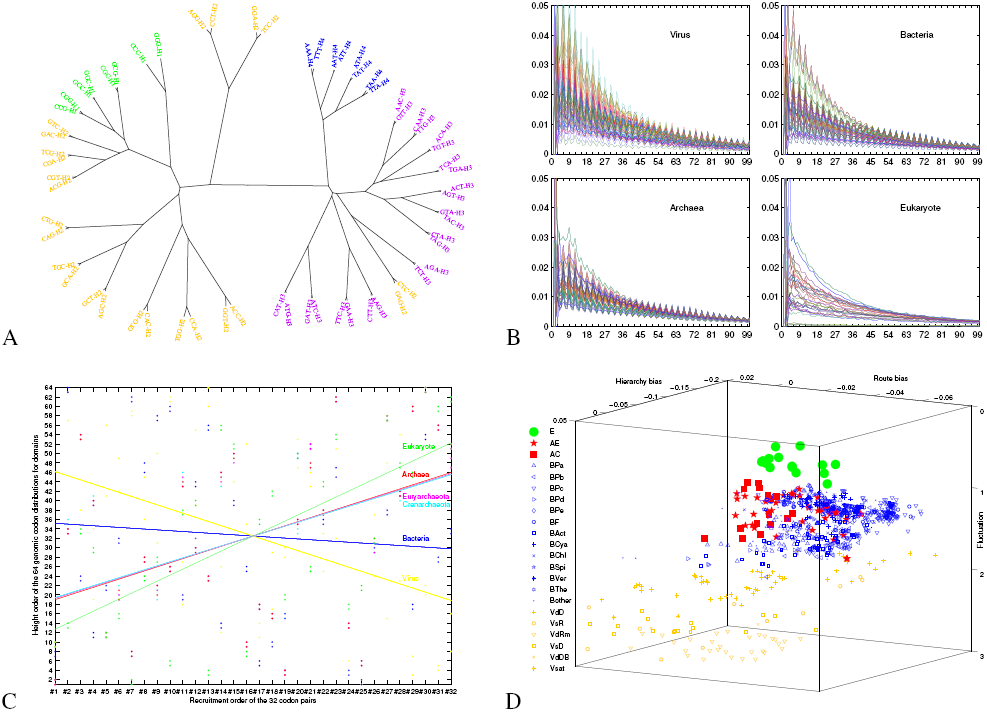
Explanation of the origin of the three domains of life. **A**, The tree of codons based on the genomic codon distributions, which agrees with the differentiation of the four hierarchies *Hierarchy* 1 − 4 (H1-4) in the roadmap. **B**, Features of genomic codon distributions for domains based on complete genome sequences. **C**, On the origin order of the domains. **D**, Classification of Virus (yellow), Bacteria (blue), Archaea (red) and Eukarya (green) in the evolution space, which is based on the roadmap Fig 1A and the complete genome sequences.

The genomic codon distribution is an essential property of a species at the sequence level. It is shown that the initial features of genomic codon distributions resulted from the evolution of the genetic code (Fig S3a1), and that the features of genomic codon distributions are relatively invariant for the large scale genome duplications in genome size evolution [21] (Fig S3a4). The features of genomic codon distributions formed during the evolution of the genetic code have been conserved throughout the history of life. Hence, both the tree of codons and the tree of species have been dug out from the genomic codon distributions of contemporary species.

The features of genomic codon distributions differ among Virus, Bacteria, Archaea and Eukarya (Fig 3B, S3b1, S3b2, S3c1). The amplitudes of the genomic codon distributions are higher for Virus, middle for Bacteria and Archaea, and lower for Eukarya (Fig 3B). The orders of the 64 average heights of the genomic codon distributions of codons differ among the domains: the height orders for Virus and Bacteria are about opposite to that for the Archaea and Eukarya (Fig S3c1). The genomic codon distribution for Eukarya is quite different from those of other domains, due to its more complicated fluctuations.

The features of genomic codon distributions can be simulated based on the roadmap (Fig S3b3, S3c2). The simulations of these statistic features at the sequence level indicate that the three domains of life originated at different stages in the evolution of the genetic code (see Appendix). The origin order of domains has been deduced from the recruitment order of codons and the average heights of the genomic codon distributions (Fig 3C).

An evolution space is constructed based on the roadmap and the features of genomic codon distributions, which helps to explain the origin of the three domains (Fig 3D). The three coordinates of the evolution space, i. e. fluctuation, route bias and hierarchy bias, are defined respectively according to the differences of genomic codon distributions between certain groups of codons in the roadmap (see Appendix) (Fig S3d2). Virus, Bacteria, Archaea and Eukarya gather together in clusters separately in the evolution space (Fig 3D). It is shown that Virus and Eukarya can be discerned by the coordinate fluctuation (Fig S3d4); Bacteria and Archaea can be distinguished by the other two coordinates (Fig 3D), where the route bias plays a primary role (Fig S3d5). Crenarchaeota and Euryarchaeota cannot be distinguished obviously in the evolution space, but they can be distinguished by a similarly defined leaf node bias (Fig S3d6, S3d7).

Another distance matrix for species has been defined according to the Euclidean distances among species in the evolution space (see Appendix). Thus a tree of life has been obtained by the evolution space (Fig S3d8), especially for eukaryotes (Fig S3d10). According to the average Euclidean distances among taxa, the evolutionary tree for Virus, Bacteria, Crenarchaeota, Euryarchaeota and Eukarya is obtained (Fig S3d9), which agrees with the three-domain tree [8] rather than the eocyte tree [9]. Actually, the features of genomic codon distributions between Crenarchaeota and Euryarchaeota are more similar than comparing them with Eukarya (Fig S3b1, 3B).

The tree of life is fundamentally determined by the evolution of the genetic code. The diversity of the genetic code itself led to the diversity of life. Major branches of the tree of life are determined by the major route bias and hierarchy bias, while minor branches of the tree of life by the minor leaf node bias and so forth. Biodiversity is intrinsic for the chiral living system, which existed at the root of the tree of life. Thus the last universal common ancestor becomes an unnecessary notion in theory. None single species can persistently exist in the non-chiral surroundings, but an ecosystem can survive owing to limited life spans of diverse individuals. Biodiversity played a crucial role in the adaptation of life to the changing Earth system, which will be explained in the next section.

## 3 Reconstruction of the Phanerozoic biodiversity curve based on genomic, climatic and eustatic data

The history of life can be studied at the species level as well as at the sequence level. Genome size is the most coarse-grained representation of a species at the sequence level. The growth trend of biodiversity at the species level can be explained in terms of statistical analysis of genomes sizes. The genome sizes of contemporary species tend to follow logarithmic normal distribution [22] [23] (Fig S4a1), which can be simulated by a statistical model based on genome duplications (Fig S4a2). According to the model, the genome size of the ancestor of a taxon is estimated by subtracting the standard deviation from the logarithmic mean of genome sizes of species in the taxon (Fig S4a7, S4a2, S4a4). By considering the origin times of taxa (Fig S4a7), the trend of genome size evolution is obtained (see Appendix) (Fig 4A). Besides, the history of Phanerozoic biodiversity is recorded in the fossils [10] [24] [25]. The biodiversity curve through the Phanerozoic [10] can be split into two parts: the exponential growth trend [26] and the biodiversity net fluctuations (Fig S4a8). It is interesting to find that the exponential growth rate of the Phanerozoic biodiversity curve is about equal to the exponential growth rate of genome size evolution (see Appendix) (Fig 4A). This indicates that the Phanerozoic biodiversity at the species level was boosted by the genome size evolution at the sequence level.

**Fig 4.**
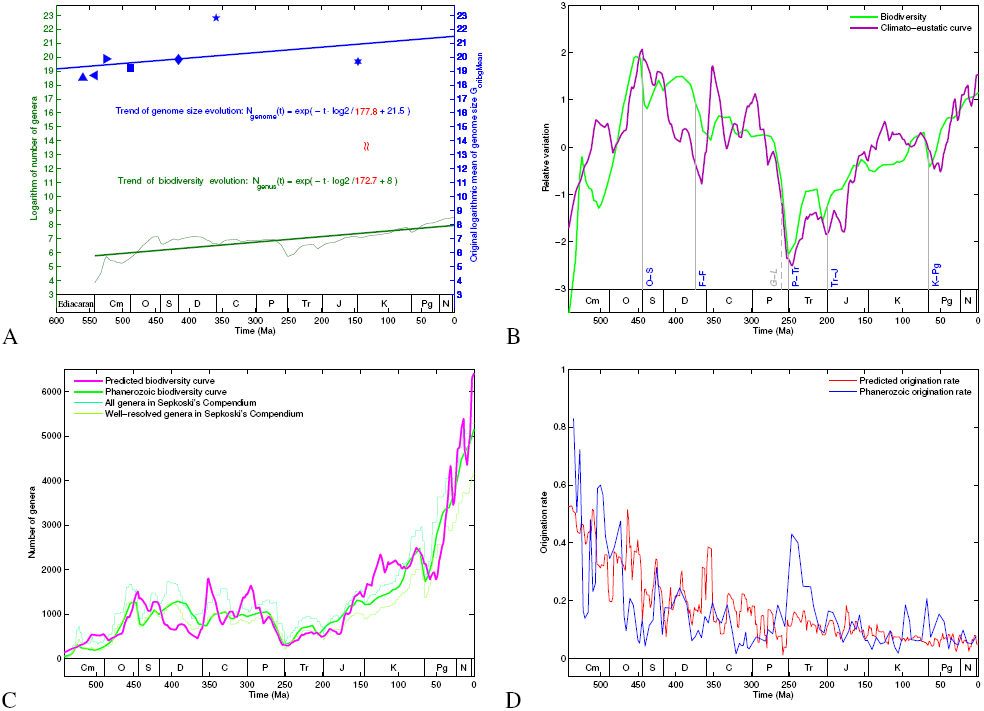
Explanation of the Phanerozoic biodiversity curve. **A**, Explanation of the trend of the Phanerozoic biodiversity curve by the trend of genome size evolution. **B**, Explanation of the fluctuations of the Phanerozoic biodiversity curve based on climate changes and sea level fluctuations, which indicates the tectonic cause of the mass extinctions. **C**, Reconstruction of the Phanerozoic biodiversity curve based on genomic, climatic and eustatic data, whose result agrees with the biodiversity curve based on fossil records. **D**, Explanation of the declining origination rate through the Phanerozoic.

The sea level fluctuations and climate changes have directly influenced the Phanerozoic biodiversity. Despite considerable differences among the following results on the Phanerozoic climate: climate indicators [27] [28] (Fig S4b1), Berner’s *CO*_2_ [29], marine ^87^*S r/*^86^*S r* [30] [31], and marine *δ*^18^*O* [31] [32], all these four methods are reasonable from their own aspects. A pragmatic approach is to average the four results (after nondimensionalization), and to take the consensus result as the climate curve in the Phanerozoic (Fig S4b2). The lower the global temperature was, the higher the global latitudinal climate gradient was (Fig S4b2). Higher climate gradient leads to more biodiversity, due to a richer environment on Earth. The eustatic curve in the Phanerozoic has been obtained by averaging the Haq’s curve [33] [34] and Hallam’s curve [35] (Fig S4b3). Hence, a climato-eustatic curve is defined by averaging the climate gradient curve and the eustatic curve (Fig S4b4), which evaluates the impact on biodiversity from Earth’s changing environments. It is shown that the biodiversity net fluctuations can be explained by the climato-eustatic curve (Fig 4B).

Thus, the Phanerozoic biodiversity curve can be reconstructed by assembling both the climato-eustatic curve and the exponential growth trend driven by the genome size evolution (see Appendix) (Fig S4c1). The reconstructed biodiversity curve based on genomic, climatic and eustatic data generally agrees with the Phanerozoic biodiversity curve based on fossil records, both in the fluctuations and in the trend (Fig 4C). This split and reconstruction scheme indicates that the combination of genome size evolution and interactions among Earth’s spheres did shape the Phanerozoic biodiversity curve (Fig S4b5).

The reconstructed biodiversity curve helps to explain the outline of the Phanerozoic biodiversity curve based on fossil records: the Cambrian and Ordovician radiations, the Paleozoic biodiversity plateau, the P-Tr mass extinction, and the Mesozoic and Cenozoic radiation (see Appendix) (Fig 4B, 4C). The five mass extinctions can be explained altogether by the climato-eustatic curve (see Appendix) (Fig 4B), which indicates that the mass extinctions resulted from multifactorial impacts of the interactions of Earth’s spheres rather than any single catastrophic event. The aggregation and dispersal of Pangaea resulted in the sea level fluctuations at the tectonic time scale, which consequently influenced the climate gradient changes at the tectonic time scale. Both sea level fluctuations and climate gradient changes substantially impacted on the mass extinctions (Fig S4b5). Marine regressions are among the explanations for the five mass extinctions in literatures. The mass extinctions during F-F, P-Tr, Tr-J and K-Pg occurred in the lower climate gradient periods (Fig S4b5). As for two phases of the P-Tr mass extinction [36], marine regression occurred during end-Guadalupian and low climate gradient occurred during end-Changhsingian (Fig S4b5). Most minor extinction events in the Phanerozoic [37] can also be explained by the climato-eustatic curve in detail (Fig 4B, S4b7).

The split and reconstruction scheme helps to explain the synchronously declining trend in both origination rate and extinction rate in the Phanerozoic (see Appendix) [38] [39] (Fig 4D, S4d3). The genome size evolution steadily boost the exponential growth in biodiversity, which resulted in the declining origination rate (Fig 4D) in consideration of the constraint on origination ability on Earth (Fig S4d1). The exponential growth trend of biodiversity demands the synchronicity between the origination rate and the extinction rate (Fig S4d3, S4d2). The living system creates redundant species at the sequence level, so as to gain excellent adaptation through high extinction rates at the species level. Thanks to the homochirality, living organisms are at home on Earth.

## 4 Discussions

It is worthwhile to test the above theory on the origin of homochirality of life experimentally. The chiral symmetry breaking through the winner-take-all game is suggested to be detected based on substitutions in the triplex DNAs under the periodic experimental conditions. Furthermore the universal genetic code is expected to be obtained in experiment by considering the increasing stabilities of base triplexes in the roadmap.

The three domains of life cannot be distinguished by the differences of genomic codon distributions between two arbitrarily selected groups of codons in the roadmap. This convinces that the evolution of the genetic code following the roadmap shaped the main branches of the tree of life. The clusterings of the three domains of life in the evolution space is anticipated to be verified by more genomic data.

## Acknowledgements

My warm thanks to Jinyi Li for valuable discussions. Supported by the Fundamental Research Funds for the Central Universities.

